# Accurate sex classification from neural responses to sexual stimuli

**DOI:** 10.1101/2022.01.10.473972

**Authors:** Vesa Putkinen, Sanaz Nazari-Farsani, Tomi Karjalainen, Severi Santavirta, Matthew Hudson, Kerttu Seppälä, Lihua Sun, Henry K. Karlsson, Jussi Hirvonen, Lauri Nummenmaa

## Abstract

Sex differences in brain activity evoked by sexual stimuli remain elusive despite robust evidence for stronger enjoyment of and interest towards sexual stimuli in men than in women. To test whether visual sexual stimuli evoke different brain activity patterns in men and women, we measured haemodynamic brain activity induced by visual sexual stimuli in two experiments in 91 subjects (46 males). In one experiment, the subjects viewed sexual and non-sexual film clips and dynamic annotations for nudity in the clips was used to predict their hemodynamic activity. In the second experiment, the subjects viewed sexual and non-sexual pictures in an event-related design. Males showed stronger activation than females in the visual and prefrontal cortices and dorsal attention network in both experiments. Furthermore, using multivariate pattern classification we could accurately predict the sex of the subject on the basis of the brain activity elicited by the sexual stimuli. The classification generalized across the experiments indicating that the sex differences were consistent. Eye tracking data obtained from an independent sample of subjects (N = 110) showed that men looked longer than women at the chest area of the nude female actors in the film clips. These results indicate that visual sexual stimuli evoke discernible brain activity patterns in men and women which may reflect stronger attentional engagement with sexual stimuli in men than women.

## Introduction

Identification of potential sexual partners is crucial for reproduction and species survival and relies strongly on visual cues in humans and other primates (Georgiadis & Kringelbach, 2012). Accordingly, visual sexual stimuli are rewarding and sexually arousing for humans (Wierzba et al., 2015) and they trigger autonomic and genital responses that prepare the body for sexual activity in both men and women (Chivers et al., 2010). Yet, human reproductive and sexual behaviour is markedly dimorphic presumably because females carry significantly higher metabolic costs of reproduction while men have higher capacity for conceiving offspring (Buss & Schmitt, 2019). Men have a stronger sexual motivation than women, as indicated by higher desired frequency of sexual intercourse, number of sexual partners, and more frequent masturbation and sexual fantasies (for a meta-analysis, see Baumeister et al., 2001). Accordingly, men experience stronger sexual arousal (Murnen & Stockton, 1997) and positive affect (Peterson & Janssen, 2007; Sarlo & Buodo, 2017) than women when viewing sexual stimuli depicting the preferred sex. Finally, men also consume erotica and pornography more than women (Rissel et al., 2017; Solano et al., 2020) and begin doing so at an earlier age (Hald, 2006). These sex differences in subjective experience and consummatory behavior suggest sexually dimorphic brain activity patterns to sexual signals. However, this issue remains hotly debated (Mitricheva et al., 2019; Poeppl et al., 2020).

Meta-analyses of neuroimaging studies show that visual sexual vs. neutral stimuli activate numerous cortical and subcortical regions spanning the emotion and reward circuit such as the brainstem, ventral striatum, amygdala, hypothalamus, thalamus, insula, cingulate, premotor cortex, and visual cortices (Mitricheva et al., 2019; Poeppl et al., 2016). Most of these studies have been conducted in men, and the few studies comparing men and women have yielded mixed results. Early small-scale studies reported stronger activation in men compared to women in hypothalamus (Hamann et al., 2004; Karama et al., 2002), amygdala (Hamann et al., 2004) and extrastriatal visual cortex (Sabatinelli et al., 2004) in response to erotic stimuli. More recent studies have found stronger activity in men in the thalamus, anterior and middle cingulate gyrus and occipital and parietal cortices (Wehrum et al., 2013) as well as in nucleus accumbens (NAc) in response to erotic pictures (Wehrum-Osinsky et al., 2014). Other studies, however, have found no evidence for sex differences (Gillath & Canterberry, 2012; Stark et al., 2005). Meta-analyses have also yielded conflicting results regarding sex differences in the processing of visual sexual stimuli: Poeppl et al (2016) found stronger engagement of the basal ganglia and hypothalamus in men whereas Mitricheva et al. (2019) found no evidence for sex differences. Thus, despite the marked sex differences in sexual behavior, the putative sex differences in the neural processing of visual sexual stimuli have remained elusive.

So far, all fMRI studies on sex differences in the processing of visual sexual stimuli have employed conventional mass-univariate general linear model (GLM) analysis. In contrast to the GLM approach, multivariate pattern analysis (MVPA) of fMRI data uses machine learning algorithms to capture activity patterns distributed across voxels that differentiate conditions or classes such as stimulus categories (Haxby et al., 2001), emotional and cognitive states (Putkinen et al., 2021; Saarimäki et al., 2015) and subject groups (Du et al., 2012; Shimizu et al., 2015). Since MVPA exploits information aggregated across voxels, this analysis approach is sensitive to spatially distributed group differences that are not detectable by conventional univariate analyses (Nummenmaa & Saarimäki, 2017). Accordingly, previous fMRI studies have successfully employed MVPA to reveal sex differences in resting state activity (Weis et al., 2020) as well as language-related tasks (Ahrens et al., 2014; Xu et al., 2020). Thus, even though univariate (voxel-wise) GLM analysis of brain responses to visual sexual stimuli may not reliably differentiate males and females, multivariate brain activation patterns induced by such stimuli could be more sensitive to such sex differences.

### The current study

Here we used multivariate pattern analysis in a large sample of subjects (N = 91, 46 males) to test whether males and females show different neural responses to sexual stimuli. In two experiments, subjects viewed sexual and non-sexual film clips and pictures while their brain activity was measured with fMRI. We found that males showed stronger activation than females particularly in the visual and prefrontal cortices in both experiments. Furthermore, using MVPA we could accurately predict the sex of the subject on the basis of the brain activity elicited by the sexual stimuli. The classification generalized across the experiments so that sex could be predicted when the classifier was trained on the data from one experiment and tested on the data from the other indicating that the sex differences were consistent across the experiments.

## Methods

### Subjects

Altogether 91 volunteers participated in the study. Four additional participants were scanned but excluded because of a neurological abnormalities revealed in their MRIs or unusable data due to a broken gradient coil. The final sample consisted of 46 males (mean age 31, range 20-57) and 45 females (mean age 32, range 20-56). All subjects reported either heterosexual (N=81, 38 females) or bisexual (N = 10, 7 females) as their sexual orientation. The exclusion criteria included a history of neurological or psychiatric disorders, alcohol and substance abuse, current use of medication affecting the central nervous system and the standard MRI exclusion criteria. All subjects gave an informed, written consent and were compensated for their participation. The ethics board of the Hospital District of Southwest Finland had approved the protocol and the study was conducted in accordance with the Declaration of Helsinki.

### fMRI Study design

To map the brain regions encoding sexual content in films, we used a paradigm (Karjalainen et al., 2017, 2019; Lahnakoski et al., 2012) in which the subjects were shown 96 movie clips with variable content (total duration 20 minutes). Nine movie clips (mean duration 11.9 s) were extracted from commercially available pornographic films and depicted female-male couples engaged in sexual intercourse. The other film clips (mean duration 17.8 s) were extracted from mainstream English language feature films and contained scenes with non-sexual human interaction (friendly discussion, arguing, violence), human actions not directly related to social interaction (e.g., eating, walking, driving), as well as scenes depicting non-human animals, landscapes and objects (Figure 1a).

**Figure 1.** Experimental design and sample stimuli for the movie (a) and picture experiment (b).

The movie clips were presented in a fixed order without breaks. Dynamic ratings with a 4-s temporal resolution were obtained for the presence of sexual content (continuous scale from 0 to 100) in the film clips from a separate sample of subjects (n=6) not participating in the fMRI study. The dynamics ratings for sexual content were averaged across subjects and then used as a regressor in the first level GLM analysis (see below).

In the second fMRI experiment the subjects were shown 300 non-erotic images from Nencki Affective Picture System (NAPS) and 24 erotic images from Nencki Affective Picture System Erotic Subset (NAPS-ERO) data sets (Marchewka et al., 2014; Wierzba et al., 2015) in a random order (total duration, 22 min) (Figure 1b). The erotic pictures depicted female-male, and female-female couples engaged in sexual intercourse or interaction. The non-erotic pictures depicted people engaged in non-sexual activities (e.g., sports) as well as animals, landscapes, and inanimate objects. Each picture was presented for 1.5 seconds followed by a fixation cross for 2-3 seconds. The data from this experiment were used to test the generalization of the multivariate pattern classification (see below). All film clips and pictures were presented via NordicNeuroLab VisualSystem binocular display. For the film experiment, audio was presented binaurally via MRI-compatible headphones (Sensimetrics S14) at a comfortable level adjusted individually for each participant.

### MRI data acquisition

The MRI data were acquired using a Phillips Ingenuity TF PET/MR 3T whole-body scanner. High-resolution (1 mm^3^) structural images were obtained with a T1-weighted sequence (TR 9.8 ms, TE 4.6 ms, flip angle 7°, 250 mm FOV, 256 × 256 reconstruction matrix). 472 functional volumes were acquired with a T2*-weighted echo-planar imaging sequence (TR = 2600 ms, TE = 30 ms, 75° flip angle, 240 mm FOV, 80 × 80 reconstruction matrix, 62.5 kHz bandwidth, 3.0 mm slice thickness, 45 interleaved slices acquired in ascending order without gaps).

### Structural and functional MRI data preprocessing

MRI data were preprocessed using fMRIPprep 1.3.0.2 (Esteban, Markiewicz, et al. 2018). The following preprocessing was performed on the anatomical T1-weighted (T1w) reference image: correction for intensity non-uniformity, skull-stripping, brain surface reconstruction, spatial normalization to the ICBM 152 Nonlinear Asymmetrical template version 2009c (Fonov et al. 2009) using nonlinear registration with antsRegistration (ANTs 2.2.0) and brain tissue segmentation. The following preprocessing was performed on the functional data: co-registration to the T1w reference, slice-time correction, spatial smoothing with an 6mm Gaussian kernel, automatic removal of motion artifacts using ICA-AROMA (non-aggressive) (Pruim et al. 2015) and resampling to the MNI152NLin2009cAsym standard space.

### Full-volume GLM data analysis

The fMRI data were analyzed in SPM12 (Wellcome Trust Center for Imaging, London, UK, (http://www.fil.ion.ucl.ac.uk/spm). To reveal regions activated by sexual content in the films, a general linear model (GLM) was fit to the data where the mean sexual content ratings were used as a regressor. For the picture experiment, the erotic and landscape pictures were modelled separately with stick functions. For each subject, contrast images were generated for the main effect of sexual content in the films and for the erotic-minus-control contrast in the picture experiment and subjected to a second-level analysis where the contrast images were compared between men and women separately for the two experiments. Clusters surviving family wise error (FWE) correction (P < 0.05) are reported.

### Multivariate pattern classification analysis

For the main multi-variate pattern analysis (MVPA), the subject-wise beta maps obtained for the sexual stimuli in both experiments were used in classifying subjects’ sex. Only gray matter voxels (n=189114) were included in the analysis. The beta maps were normalized to have zero mean and unit variance before the application of MVPA. For all analyses except for the cross-classification between experiments (see below), a linear support vector machine (SVM) implemented in Python was used in the classification due to its ability to perform well in high-dimensional spaces. To test the performance of the classifier, leave-one-subject-out cross-validation was performed, where the classifier was trained on data from all except one subject and tested on the hold-out subject data. Classification accuracy was defined as the proportion of correctly classified subjects. To determine whether the classification accuracy was above the theoretical 50% chance level, we generated a null distribution for the classification accuracy by repeating the analysis 1000 times while shuffling the category (male / female) labels. Classification accuracy was determined significant if it was higher that 95% of the accuracies obtained with the randomly shuffled labels.

We also tested the generalizability of the sex classification by training the classifier on the film data and testing it with the data from in the pictures experiment (contrast erotic pictures > landscapes) and vice versa. This analysis was otherwise identical to the main MVPA described above, except that in the leave-one-subject-out cross-validation the test data was taken from the pictures experiment if the classifier was trained on the film data, and from the film experiment if the classifier was trained on the pictures data. For the cross-classification between the film and picture experiment, a non-linear kernel was employed as this produced higher classification accuracies than the linear kernel.

Finally, we tested whether successful sex classification could be achieved also with non-sexual stimuli, and whether the sex-specific activation patterns would generalize across sexual and non-sexual stimuli. Specifically, the subjects rated the presence of non-sexual human actions involving vocalizations (e.g., neutral discussion, laughter, arguing) in the films and the subject-wise beta maps obtained for this control dimension were subjected to an identical MVPAs as those obtained for the sexual content in the film clips. Cross-classification was performed between the control dimension and responses to the sexual films and the sexual pictures.

### Eye tracking

To test for sex differences in attention allocation while viewing the sexual stimuli, we conducted an eye movement experiment. A total of 110 subjects (43 males, mean age 27 years) not participating in fMRI volunteered for the study. Subjects viewed a subset of the videos presented in the fMRI experiment while their eye movements were recorded using Eyelink 1000 eye-tracker (SR Research, Mississauga, Ontario, Canada; sampling rate 1000 Hz, spatial accuracy better than 0.5°, with a 0.01° resolution in the pupil-tracking mode). A nine-point calibration and validation were completed at the beginning of the experiment and then after 1/3 and 2/3 of the experiment had elapsed. Saccade detection was performed using a velocity threshold of 30°/s and an acceleration threshold of 4000°/s2. Dynamic regions of interest (ROIs) were drawn frame-by frame on the face, chest, genital, and buttocks areas of the male and female characters in the sexual scenes; the remaining clips were not analyzed in this study. We computed the clip-wise proportional dwell time (%) for each region of interest and analyzed the effects of ROI as well as actor and subject sex on the dwell times using linear mixed model (LMM).

### Questionnaires

The fMRI study participants filled an online questionnaire about positive and negative emotions induced by pornography (full ratings were obtained from 86 participants, 44 males). Specifically, the subjects rated on a scale of 1-10 how strongly they felt sexual arousal, joy, curiosity, disgust, shame, sadness and anger while viewing sexually explicit media. The subjects also reported how often they viewed pornography (never, less than once a month, less than once a week, once a week or more, daily, several times a day). Subjects also rated on how often they engaged or wanted to engage in kissing, masturbation, vaginal or anal intercourse, oral sex and sexual fantasies on a scale ranging from never to several times a day (Derogatis & Melisaratos, 1979). Summary scores were computed for the actual and desired frequency of these sexual behaviors.

## Results

### Emotions elicited by sexual stimuli in males and females

Both males and females reported high positive emotions and low negative emotions towards pornography and moderate-to-high levels of pornography consumption (**Figure 2a**). Male participants reported stronger sexual arousal (*t*(85) = 2.01, *p* < .05) and joy (*t*(85) = 3.63, *p* < .001) and less shame (*t*(85) = 2.83, *p* < .01) than females. Males also consumed pornography more frequently than females: the most typical answer for males was at least once a week while the most answer for females was at least once a month (X^2^ (5, N = 86) = 25.5, p < .000) (**Figure 2c**). The frequency of pornogpraphy consumption correlated positively with sexual arousal (*r* = .44) and joy (*r* = .34) and negatively with shame (*r* = -0.24) (Supplementary Figure 4). Men also reported higher actual (*t*(83.55) = 2.00, *p* < .05) and desired (*t*(83.55) = 2.53, *p* < .05) frequency of sexual activities (**Figure 2b**)

**Figure 2.**
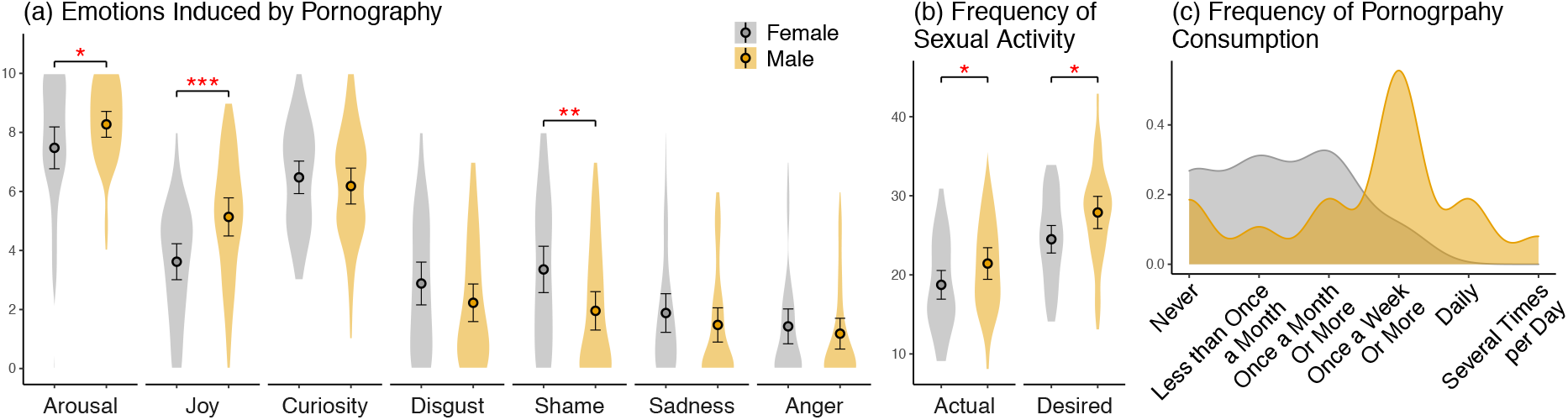
(a) Emotions induced by pornography in male and female participants. (b) The actual and desired frequency of various sexual activities. (c) Frequency of pornography consumption in males and females. * *p* < .05 ** *p* < .01, *** *p* < .001

### Brain responses to sexual stimuli

Modeling the BOLD data for the movie experiment with the sexual content regressor revealed activity in visual areas in the lateral occipital cortex extending to the inferior temporal and fusiform gyri. There was also extensive activation of regions associated with emotion and reward such as the amygdala, brainstem, thalamus, ventral striatum, insula, ACC and OFC. Activation was also observed in sensorimotor regions in the precentral gyrus, SMA and cerebellum (**Figure 3a**). Viewing erotic images in the picture experiment activated a largely overlapping network of cortical and subcortical regions (**Figure 3b**).

**Figure 3.**
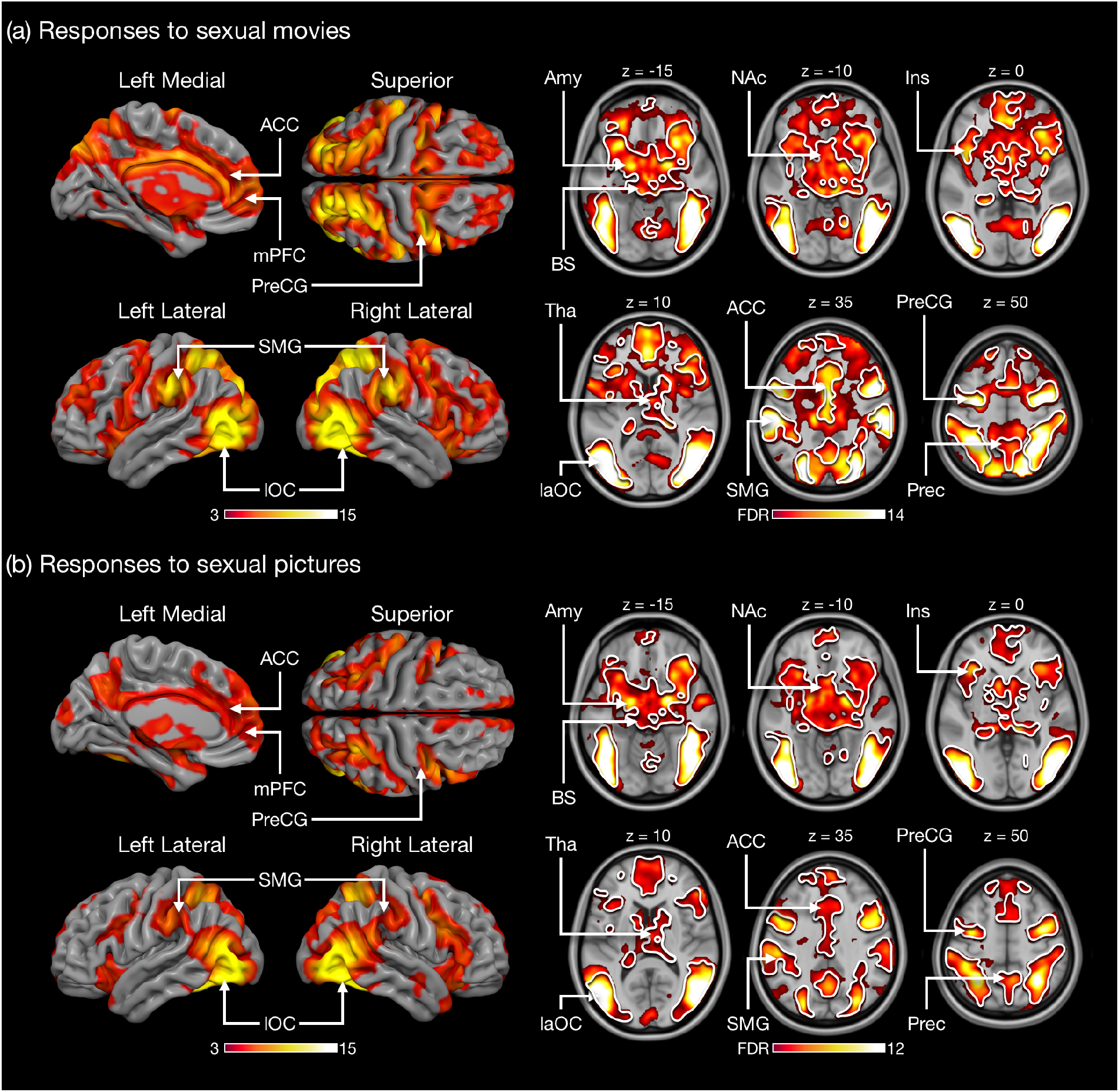
Brain regions responding to sexual content in the film clips (a) and pictures (b) (thresholded at *P* < 0.05, FWE corrected at cluster level). The white outline represents the overlap in activation across the movie and picture experiments. The colourbars represents the *t*-value. Amy = amygdala, ACC = anterior cingulate cortex, Ins = insula, laOC = lateral occipital cortex, mPFC = medial prefronal cortex, NaC = nucleus accumbens, SMG = supramarginal gyrus, Tha = thalamus, Prec = precuneus, PreCG = precentral gyrus.

### Sex differences

In the film experiment, males showed stronger response to sexual content compared to females in the lateral occipital cortex, occipital pole, fusiform gyrus, frontal pole, middle frontal gyrus, intraparietal sulcus (IPS) and precentral gyrus/middle frontal gyrus including the frontal eye field (FEF) (**Figure 4a, Figure 5**). In contrasts, females showed stronger activation than males in auditory cortical regions, parietal operculum and the right precentral gyrus than males in responses to sexual content in the film clips (**Figure 4a**). In the picture experiment, males showed stronger response than females for the contrast erotic pictures > landscapes in a largely the same areas which was activated more strongly in males in the movie condition (**Figure 4b, Figure 5**). Unlike in the movie experiment, no region showed stronger activity in females than in males (the results of separate follow-up analyses for the female-male and female-female picture subcategories were concordant with the main analysis, see Supplementary Figure 1).

**Figure 4.**
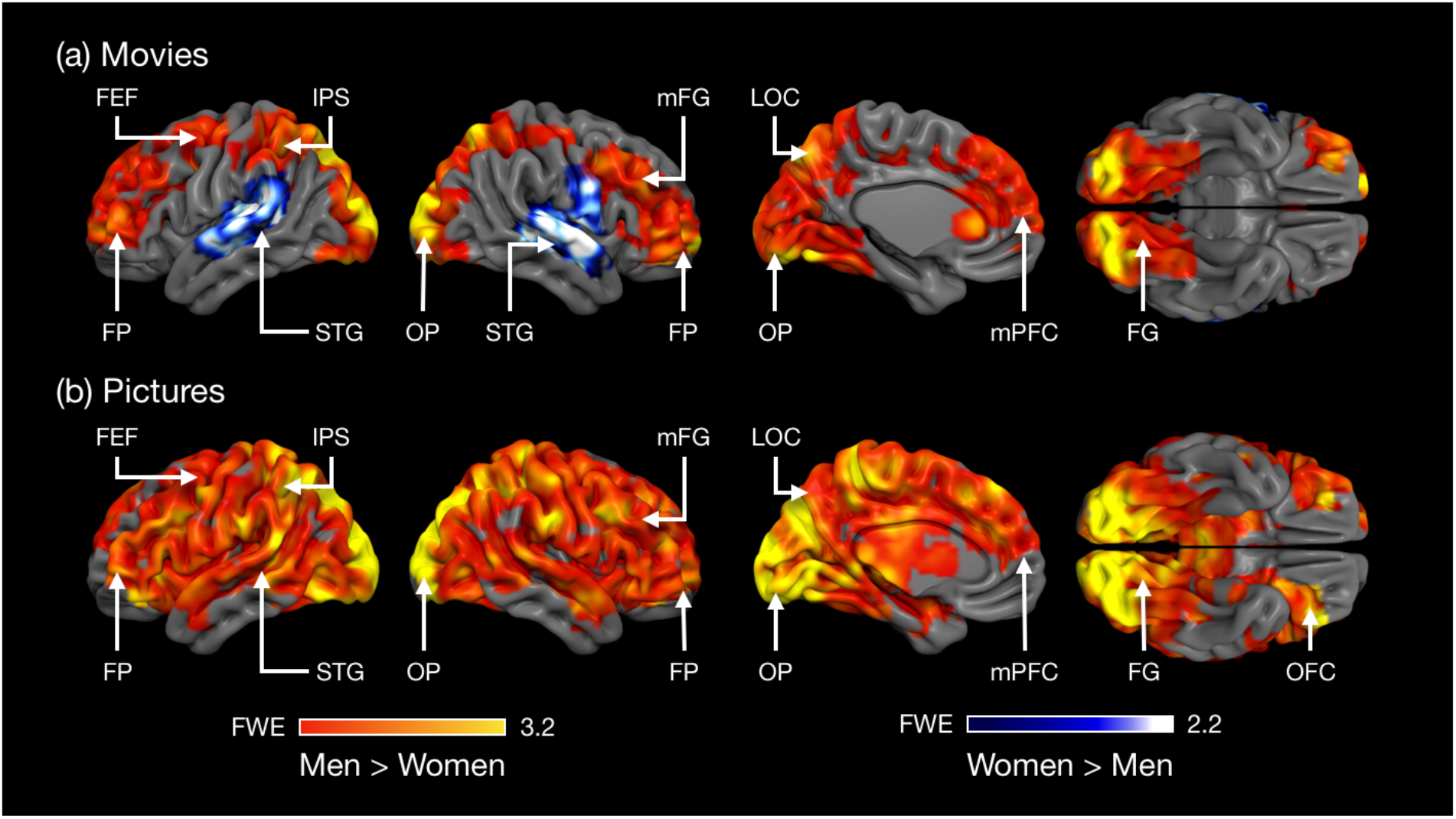
(a) Brain regions showing stronger responses to sexual content in males than in females in the movie experiment. (b) Brain regions showing stronger responses in males than in females in response to the sexual pictures. The activation maps are thresholded at *P* < 0.05, FWE corrected at cluster level. The colourbars show the *t*-value. FEF = frontal eye field, FG = fusiform gyrus, FP = frontal pole, LOC = lateral occipital cortex, mFG = medial frontal gyrus, vmPFC = ventromedial prefrontal cortex, IPS = intraparietal sulcus, OFC = orbitofrontal cortex, OP = occipital pole, SMG = supramarginal gyrus, STG = superior temporal gyrus.

**Figure 5.**
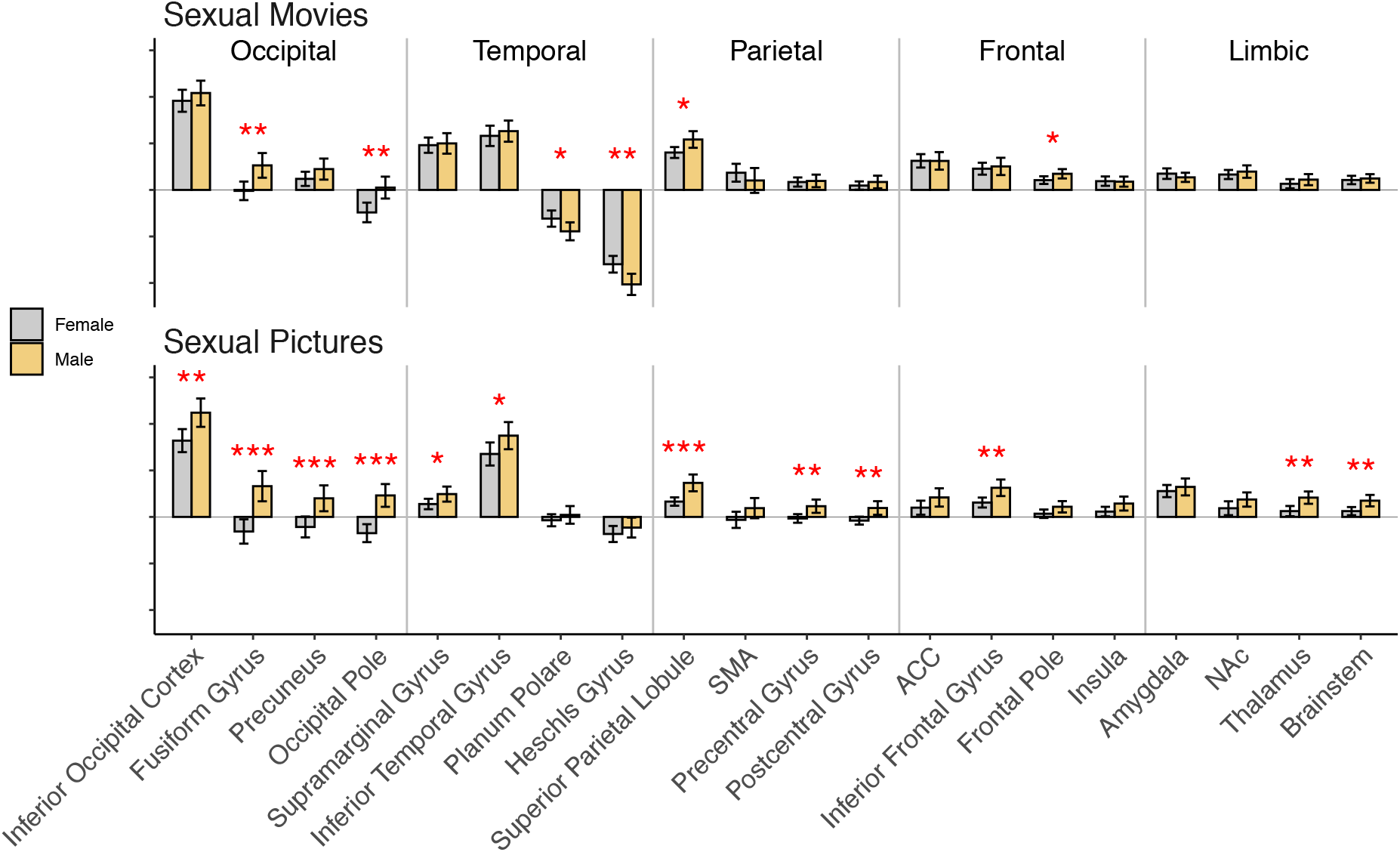
Mean beta weigths (arbitrary scale) within selected ROIs for both sexual movies and pictures. The error bars indicate 95% confidence intervals. Asterisks denote significant sex-differences at the ROI level. ∗ *p* < 0.05, ∗∗ *p* < 0.01, ∗∗∗ *p* < 0.001.

### Multivariate pattern classification

Males and females were classified above chance level based on their responses to both the erotic movie clips (mean accurasy = 76%) and pictures (mean accurasy = 64%; Figure 6). The classifier was also generalizable across the experiments: classification accuracy was above chance level even when the the classifier was trained on the Movie data and tested on the Pictures data (mean accuracy 66%) and vice versa (mean accuracy 66%). We also tested whether misclassifications could be attributed to bisexual orientation but this was the case for only one such female subject. Similarly, self-reported emotions towards pornography or sexual drive were not associated with classification accuracy (see Supplementary material).

**Figure 6.**
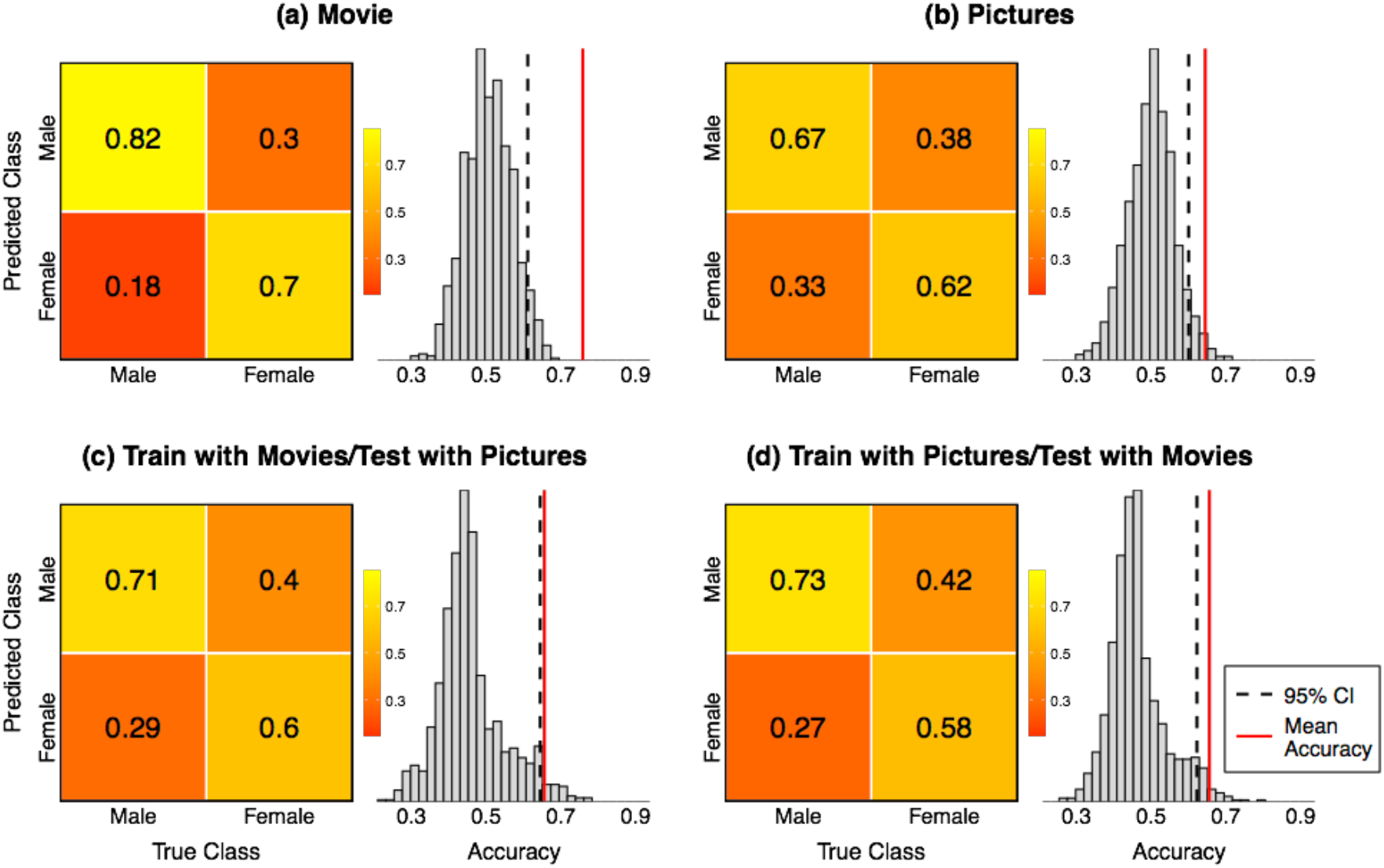
Confusion matrices and null distributions for the sex classification for the Movie and Pictures tasks and the cross-classification across these experiments. The numbers in the confusion matrices indicate the proportions of true and false predictions for males and females. The histograms show the null distribution for the classification accuracy. The red vertical lines indicates the mean classification accuracy, and the dashed vertical line the upper confidence interval limit (95% quantile) of the null distribution.

Classification was also accurate with the control dimension in the movie experiment (ie. training and testing with the respones to the control dimension, mean accuracy 68%). However, the classification failed when the responses to the control dimension was used for training and the responses to erotic clips for testing and vice versa (mean accuracies 48% and 44%, respectively, see Supplementary Figure 2) indicating that, in contrast to the cross classification with erotic movies and pictures, the classification with the control dimension did not generalize to classification with responses to the sexual film clips. This indicates the succesfull sex classification with the sexual stimuli did not reflect general sex differences in audiovisual processing but was based on processing of the sexual content in the films and pictures.

### Eye tracking results

Compared to women, men looked longer at the female actors’ chest region (F(1,107) = 11.089, *p* < 0.001). Women looked longer at males actors’ faces than men did (F(1,107) = 4.0619, *p* < 0.05). No significant sex differences were found for the other areas of interest (**Figure 7**).

**Figure 7.**
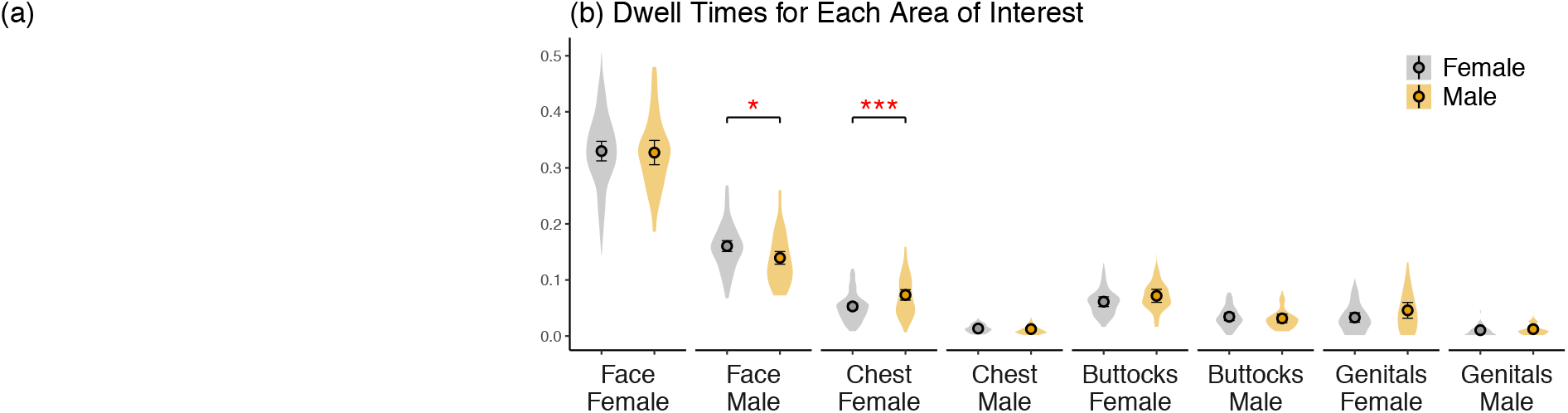
(a) Sample mean fixation distribution (heatmap) and individual fixations for the sexual movie clips (*not shown due to Biorxiv policy to avoid the inclusion of photographs of people*) (b) Average dwell times for the different areas of interests for male and female participants.

## Discussion

Our main finding was that sexual stimuli elicit discernible patterns of brain activation in men and women. Sexual movie clips and pictures elicited widespread activation across the brain in both males and females. Activations were observed in regions associated with reward and emotion (e.g. brainstem, basal ganglia, thalamus, ACC, amygdala, and medial prefrontal cortex) and in somatosensory and motor cortices (pre- and postcentral gyrus, SMA) implicated in sexual arousal (Georgiadis & Kringelbach, 2012). Activations were also observed in visual regions in the occipital and inferior temporal cortices and in the dorsal attention network (frontal eye fields, FEF and intraparietal sulcus, IPS). Men showed on average stronger responses than women particularly in visual regions in the occipital cortex and fusiform gyri, in the dorsal attention network as well as in various prefrontal regions. Notably, using multivariate pattern classification we were able to accurately predict the sex of the individual subjects. The classifier generalized across the movie and picture experiments, underlining the consistency of the sex-specific response patterns. These results indicate that, although visual sexual stimuli engage similar networks in men and women, brain activity patterns induced by such stimuli are different across sexes.

### Brain activity patterns induced by visual sexual stimuli predict individuals’ sex

Using multivariate pattern classification of the brain responses to the sexual movies and pictures, we were able to accurately classify the subjects as males or females. This indicates robust sex differences in the brain responses to sexual signals. To our knowledge, there are no previous studies employing sex-classification with visual sexual stimuli, but the classification accuracies achieved in the current study are comparable to those obtained in sex classification with resting state fMRI (Satterthwaite et al., 2015; Weis et al., 2020; C. Zhang et al., 2018; X. Zhang et al., 2020) and functional connectivity during a semantic decision task (Xu et al., 2020). The accuracy was better in the movie (76%) than in the picture experiment (66%). This likely reflects the fact that audiovisual movies are more representative of the natural environment, and consequently activate the brain more strongly and consistently than still photos (Hasson et al., 2010). Importantly, above chance level classification was achieved even with cross-classification where the classifier was trained on the data from one experiment and tested on data from the other indicating that the male/female-typical brain activation patterns evoked by sexual signals were consistent across the experiments. This indicates that the sex-specific brain responses indeed reflected the processing of the sexual content shared across the dynamic videos and still pictures.

Virtually the same regions showing sex differences in the GLM analysis (see below) also contributed the most to the classification with SVM in both experiments as indexed by the high correlation between the SVM weights and the beta values for the sex difference 2nd level contrasts (*r* = .7 for the movie experiment and *r* = .6 for picture experiment). Namely, occipital cortex and fusiform gyrus and frontal regions showed strong weights indicative of male category while temporal regions showed the strong voxel weights indicative of female category (see supplementary Figure 3). Interestingly, above chance level sex classification was obtained even with the responses to the control dimension. However, the cross-classification between the responses to the control dimension and the sexual stimuli was at chance level indicating that different activity patterns contributed to the sex classification for sexual vs non-sexual stimuli. In line with this, the spatial distribution of the SVM weight across the brain for the sexual stimuli and the control dimension were dissimilar as illustrated by the low correlation between the voxel weight maps for the sexual stimuli and the control dimension (*r* = -.17 for the movie experiment and *r* = .07 for the pictures experiment).

Although the brain activity patterns evoked by visual sexual stimuli were predictive of subject sex, some subjects were misclassified demonstrating that these brain activity patterns were not fully sexually dimorphic (cf. Joel & Fausto-Sterling, 2016). Some previous sex classification studies suggest that misclassification is associated with a characteristic cognitive or affective profile (Satterthwaite et al., 2015; Zhang et al., 2021). In the current study, the misclassified males reported lower negative emotions towards pornography than the correctly classified males, but no other differences were found between the correctly and incorrectly classified subjects (see supplementary material). Interestingly, those subjects who identified themselves as bisexual were no more likely to be misclassified than those who identified themselves heterosexual suggesting that brain activity patterns evoked by sexual stimuli are not dependent on bisexual vs. heterosexual orientation. It is possible that factors such as sexual history or more nuanced sexual preferences may contribute to the misclassification.

### Sex-dependent activation of visual and attentional circuits

Sexual stimuli activated occipitotemporal visual regions consistently in both experiments, suggesting attentional modulation of visual cortical activity for sexually salient stimuli. Event-related potential studies indicate that human bodies with visible (versus hidden) sexual signals induce amplified responses already <200 ms from stimulus onset demonstrating facilitated processing of visual sexual cues early in visual processing stream (Alho et al., 2015; Hietanen et al., 2014; Hietanen & Nummenmaa, 2011). Occipital activity was particularly strong in the putative body sensitive regions (the ‘extrastriate body area’) in the lateral occipital cortex (Downing et al., 2001) suggesting amplified processing of sexual information in the human body recognition systems (e.g. Ponseti et al., 2006). Men showed stronger activity than women in a cluster extended from V2 along the fusiform gyrus (cf. Sabatinelli et al., 2004; Wehrum et al., 2013) which suggests that sexual stimuli trigger stronger attentional amplification of visual cortical activity in men than in women. Heightened attention towards visual sexual cues facilitates sexual arousal (Dawson & Chivers, 2016) which may explain previous findings that the activation of occipitotemporal visual regions is positively associated with measures of penile erection and subjective sexual arousal (Arnow et al., 2002; Moulier et al., 2006).

Eye tracking studies indicate that men show an attentional bias towards the explicitly sexual aspects of visual sexual stimuli (Nummenmaa et al., 2012; Rupp & Wallen, 2007). Our control experiment with eye tracking revealed that men looked longer at the chest area of the nude female actors in the movie clips than women did (approximately 7% vs. 5% of the video duration in men vs. women, respectively). Women, in turn, tended to look at the male actors faces slightly longer than male subjects did. These subtle sex differences in the locus of attention may partly account for the sex differences in brain activation in the visual cortices. Accordingly, there is evidence that focusing attention on the emotionally arousing aspects of visual stimuli increases activity in the visual regions in the fusiform gyrus and lateral and interior occipital cortices (Dolcos et al., 2020; Ferri et al., 2013). Both the sexual videos and pictures activated intraparietal sulcus (IPS) and frontal eye fields (FEF) which are central nodes in the dorsal attention network supporting controlled, top-down attention (Corbetta & Shulman, 2002). In both experiments, men showed stronger activation in a parietal cluster that extended to the IPS as well as in middle frontal gyrus/precentral gyrus extending to the FEF suggesting that visual sexual stimuli engage dorsal attention network more strongly in men that in women.

Women showed stronger activation than men only in the movie experiment in auditory cortical regions. This result suggests that women responded more strongly to the audio track in the sexual video clips which consisted mostly of non-verbal female vocalizations communicating sexual pleasure. A number of studies have shown that affective vocalizations, including sexual ones (Fecteau et al., 2007), elicit stronger auditory cortical activity than neutral voices (Frühholz et al., 2016). Behavioral studies suggest a slight female advantage in emotion recognition from non-verbal emotional vocalizations (Thompson & Voyer, 2014) but sex differences in affective sound processing has not been studied extensively with neuroimaging (however, see Ethofer et al., 2007). Our results tentatively suggest that females respond stronger to non-verbal sexual vocalizations and thereby that the stronger male reactivity to sexual cues might be specific to visual domain. However, as attention towards visual stimuli attenuates auditory cortical activity (Johnson & Zatorre, 2006; Molloy et al., 2015), another explanation is that this group difference reflects stronger reduction in auditory cortical activity in men due to stronger attention towards the visual sexual cues in the videos in men compared to women. In line with this interpretation both men and women showed reduced auditory cortex activity for the sexual videos as indicated by the negative beta weights in auditory cortex (**Figure 5**).

Evolutionary accounts posit that men and women have evolved different mating strategies in domains where they have faced different adaptive challenges (Buss & Schmitt, 1993). Lower obligatory parental investment in men has presumably given rise to the stronger preference for short-term mating and sexual variation in men, as these have increased the probability of genetic success more for men than for women. Men may also have evolved a preference for physical features associated with youth since such cues signal fertility and many years of potential future reproduction (Buss & Schmitt, 2019). The type of pornography consumed by men often simulates short-term sexual encounters with novel young women (Malamuth, 1996; Salmon & Diamond, 2012). Thus, men’s higher attentional engagement with sexual stimuli might reflect evolved preferences for sexual variety and physical cues of reproductive potential. Such biological biases probably interact with cultural norms in shaping sex-typical preferences as evidenced by cross-cultural variation and changes across time in the magnitude of these sex differences (Buss & Schmitt, 2019; Petersen & Hyde, 2011).

### Emotion circuit activation in men and women

Both males and females experienced strong positive emotions and only weak negative emotions towards pornography although men reported slightly higher feelings of sexual arousal and joy and less shame than women. In accordance with the incentive value of sexual stimuli, both experiments activated the limbic and mesolimbic regions associated with reward and emotion such the ventral striatum, amygdala and hypothalamus. Activity in the ventral striatum including nucleus accumbens (NAc) has been reported consistently in both men and women in studies employing visual sexual stimuli (Hamann et al., 2004; Ponseti et al., 2006; Walter et al., 2008; Wehrum et al., 2013). NAc activity is positively associated with subjective pleasure induced by visual sexual stimuli (Sabatinelli et al., 2007) and is presumably mediated by dopaminergic neurotransmission. Accordingly, NAc activity in response to visual sexual stimuli is modulated by exogenous dopamine manipulations (Oei et al., 2012) and animal studies indicate that mesolimbic dopamine release is crucial for sexual motivation (Pfaus, 2009). Similarly to NAc, amygdala and hypothalamus are involved in the processing of various types of rewards and according to a meta-analysis (Sescousse et al., 2013), these structures are even more strongly engaged by erotic stimuli than food or monetary rewards. The amygdala is considered critical for the detection of biological relevance (Pessoa, 2010; Sander et al., 2003) and prior studies have found amygdala activity in response to visual sexual stimuli in both men and women (Hamann et al., 2004; Zhu et al., 2010). The hypothalamus controls homeostasis and autonomic responses via neuroendocrine function. Hypothalamic activity is correlated with erection during exposure to visual sexual stimuli (Arnow et al., 2002; Ferretti et al., 2005) and its activity is modulated by sexual content in men and women even when emotional arousal and valence of the stimuli are controlled for (Walter et al., 2008). Unlike some previous studies, we did not observe sex differences in the activation of the amygdala (Hamann et al., 2004), hypothalamus (Karama et al., 2002) or NAc (Wehrum-Osinsky et al., 2014) suggesting that activity evoked by visual sexual stimuli in these does not reliably differentiate men and women. Overall, the sex differences observed in the GLM analysis were most consistent in cortical regions although activity in the brainstem and thalamus were stronger in men than in women in the pictures experiment.

Finally, in both experiments, the sexual stimuli activated the primary and secondary somatosensory cortices and insula in both men and women. These regions are involved in the processing bodily sensations and interoceptive feedback (Craig, 2002). Prior studies have reported secondary somatosensory (SII) activity in both men and women in response to visual sexual stimuli (Arnow et al., 2009; Ferretti et al., 2005) and SII appears to be more generally involved in perception of touch (Keysers et al., 2010). Insula activation has also been found to correlate with penile tumescence in men (Moulier et al., 2006; Mouras et al., 2008) and the temperature of external genitalia in women (Parada et al., 2018) while viewing visual erotic stimuli. We also observed activation in the ACC, which is a common finding in studies employing visual sexual stimuli (Stoléru et al., 2012) presumably because cingulate activity is coupled with autonomic arousal (Beissner et al., 2013).

## Limitations

Subject-specific emotion ratings and physiological arousal responses were not acquired from the participants in the fMRI experiment; thus we could not directly link the haemodynamic data with direct indices of sexual arousal. The majority of our subjects identified as exclusively heterosexual and thereby we were unable to test the effects of sexual preference irrespective of gender and our results may not generalize to individuals with non-heterosexual preference.

## Conclusions

We conclude that sexual stimuli elicit discernible patterns of brain activation between men and women. Visual sexual stimuli engaged regions supporting reward, bodily sensations attention, and visual processing irrespective of sex. However, sexual stimuli activated visual cortex and prefrontal regions more strongly in men than in women. Brain activity patterns induced by sexual movies and pictures contain information about the sex of the subject that allows accurate sex-classification with multivariate pattern classification. Thus, despite substantial overlap in regions activated by sexual stimuli in men and women, brain activity patterns associated with such stimuli differ between men and women.

## Supplementary

### Brain responses to pictures of female-male and female-female couples

Separate analyses for the erotic picture categories indicated that males showed stronger activity in responses to both the pictures with male-female and female-female couples (**Supplementary Figure 1**). Female subject did not show stronger activity than the males for neither of the erotic picture categories.

**Supplementary Figure 1.**
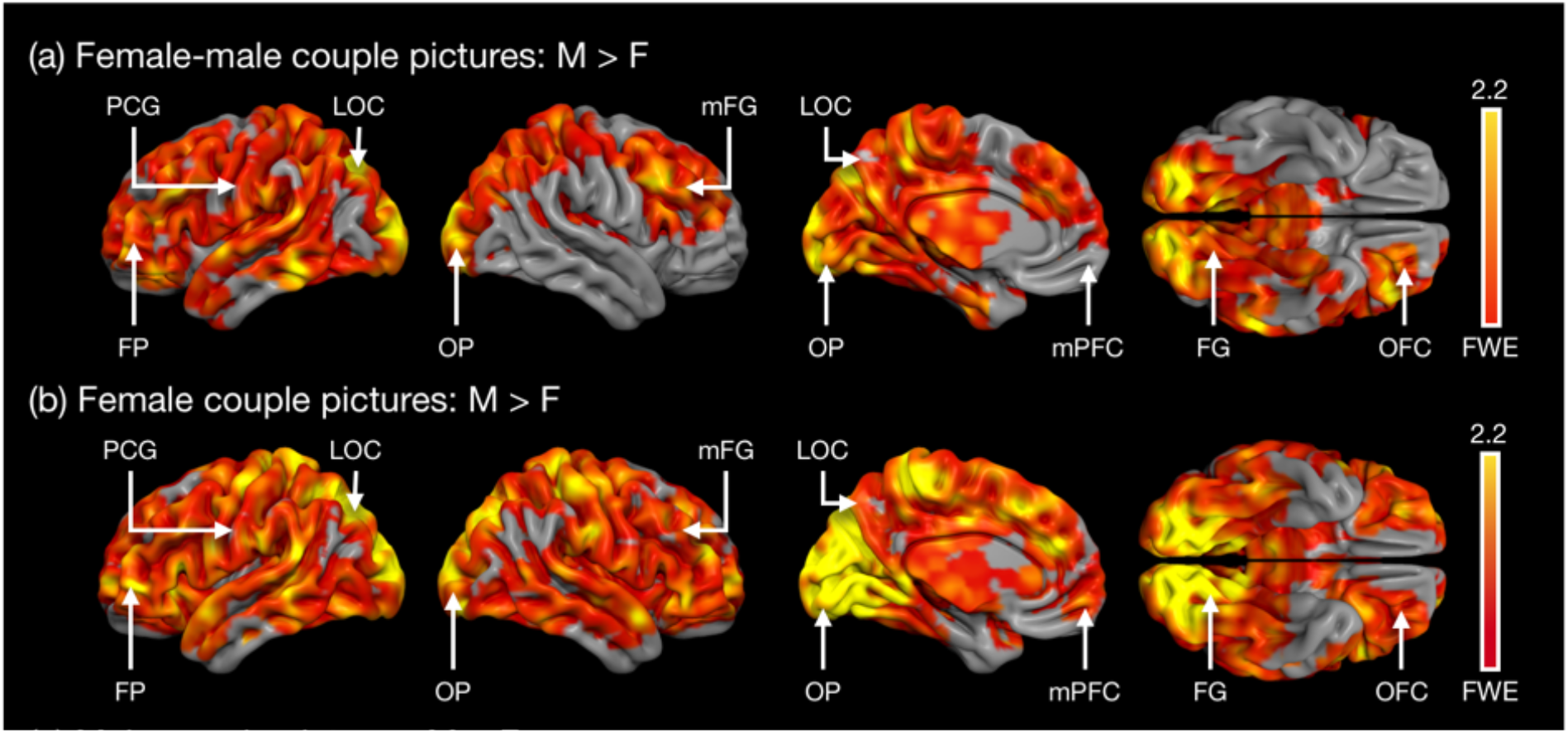
Sex differences (male > female) in response to erotic pictures depicting female-male (a), female-female (b) couples. The activation maps are thresholded at *P* < 0.05, FWE corrected at cluster level. The colourbars represents the *t*-value. FG = fusiform gyrus, FP = frontal pole, LOC = lateral occipital cortex, mFG = medial frontal gyrus, mPFC = medial prefronal cortex, NAc = nucleus accumbens, OFC = orbitofrontal cortex, OP = occipital pole, SMG = supramarginal gyrus, STG = superior temporal gyrus, PCG = precentral gyrus.

### Multivariate pattern classification: Control dimension

**Supplementary Figure 2.**
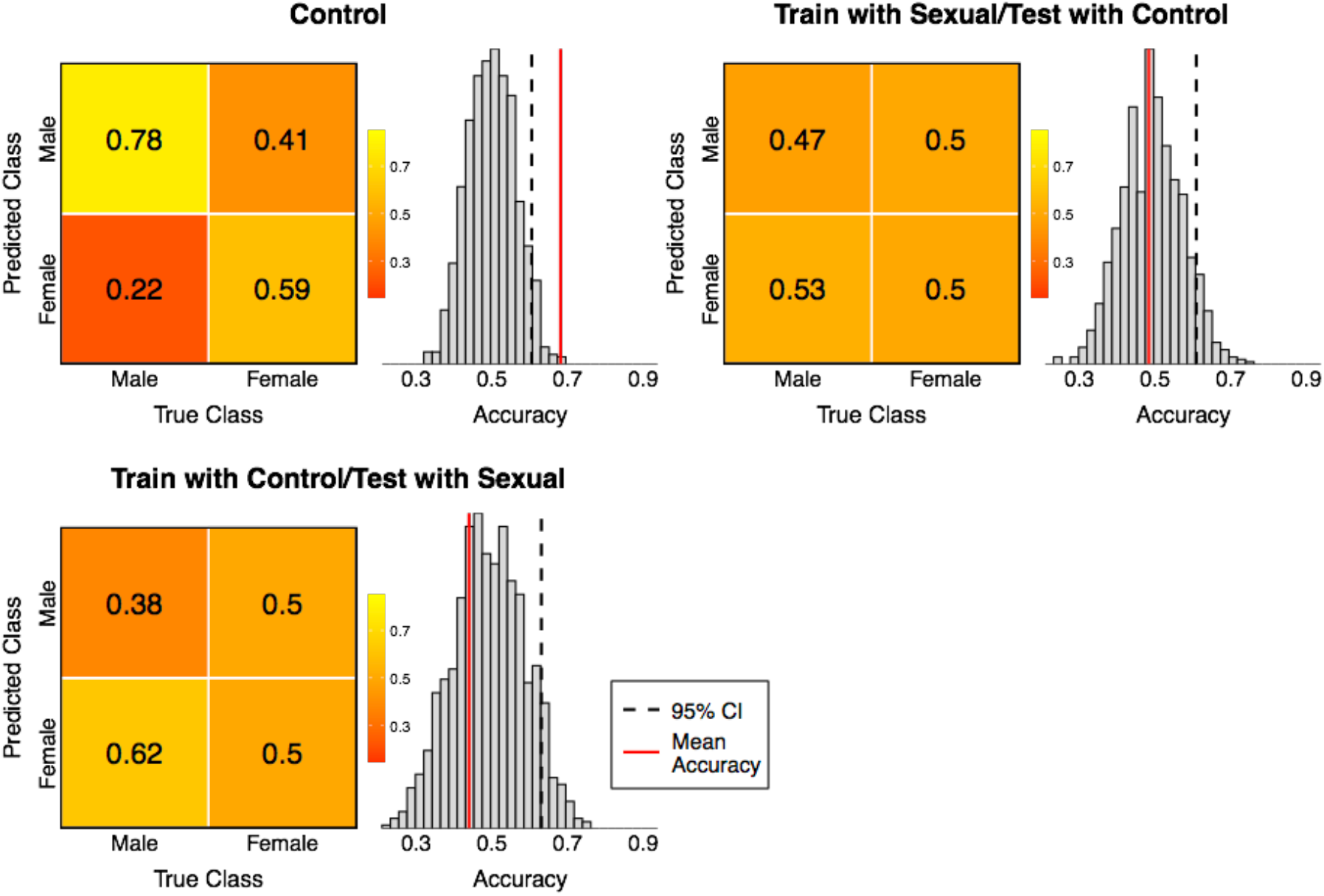
The confusion matrices and permutation results for the sex classification for the control dimension in the movie experiment and the cross-classification across the control and sexual content dimensions. The numbers in the confusion matrices indicate the proportions of true and false predictionsfor males and females. The histograms show the null distribution for the classification accuracy. The red vertical lines indicates the mean classification accuracy, and the dashed vertical line the upper confidence interval limit (95% quantile) of the null distribution.

**Supplementary Figure 3.**
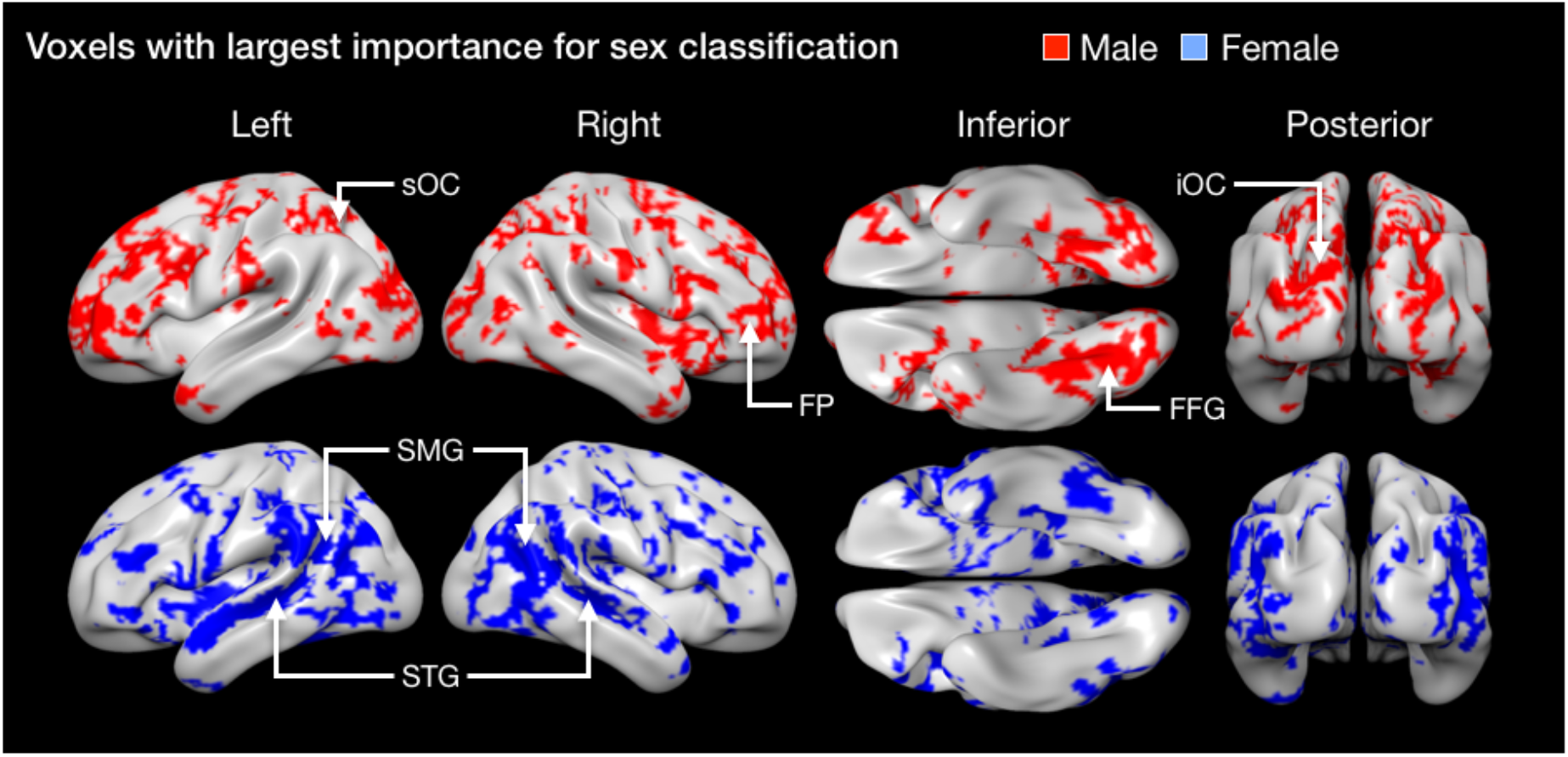
Voxels with the highest importance for the sex-classification in the movie experiment. The red regions depict voxels that were most indicative of male category and the blue regions depict voxels that were most indicative of female category. For both categories the top 40% of the voxels are shown. FP = Frontal pole, FFG = Fusiform gyrus, SMG = Supramarginal gyrus, STG = Superior temporal gyrus, iOC = inferior occipital cortex, sOC = superior occipital cortex.

### Questionnaire responses of correctly classified and misclassified subjects

We tested whether misclassified subjects would show more sex-atypical emotional responses to pornogaphy. A repeated measures analysis of variance (ANOVA) of questionnaire data revealed a significant Emotion ξ Sex ξ Correctly vs Misclassfied interaction (F(7,483) = 3.767, p < .01) on the ratings for emotion elicited by pornography. According to post-hoc pair-wise comparisons this interaction resulted from the misclassified males showing less anger, disgust, sadness and shame than the correctly classified males (all p < .05). No differences were found between the correctly vs. misclassified females. The corresponding ANOVAs on the actual and desired frequency of different sexual activities (Derogatis, 1979) did not show any differences between the correcly classified and misclassified subjects.

**Supplementary Figure 4.**
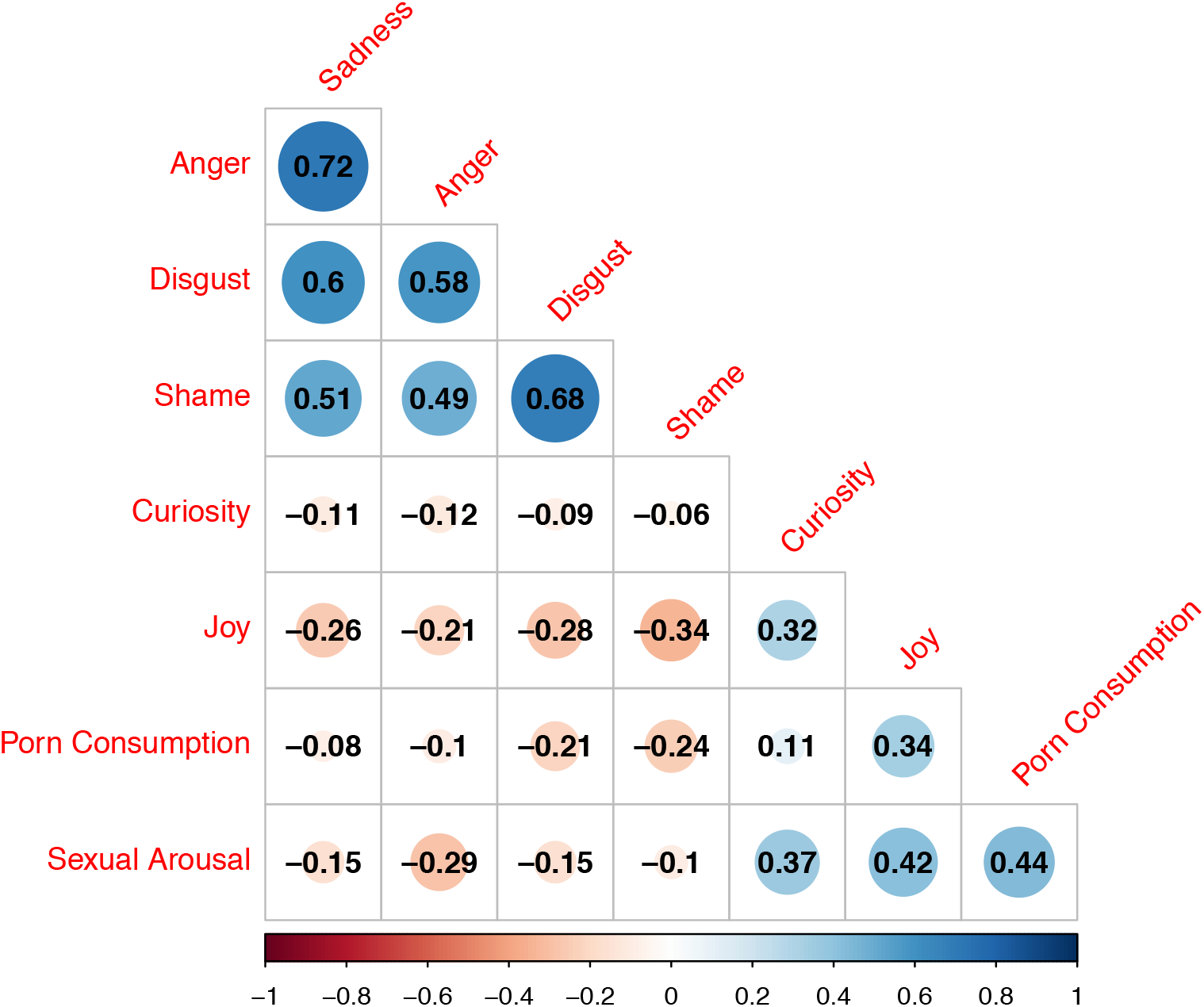
Correlations between self-ratings of emotions evoked by pornography

